# Similar weapons play different roles in bacterial competition

**DOI:** 10.64898/2026.03.03.709350

**Authors:** Sean C. Booth, Connor Sharp, Kevin R. Foster

**Author notes:** Equal Contribution (Sean C. Booth, Connor Sharp).

## Abstract

Bacteria are aggressive organisms that inhibit and kill their competitors using a diverse range of molecular weapon systems. Surprisingly, strains commonly carry multiple similar weapons that appear to perform the same function. Here we investigate the evolutionary benefits of this apparent functional duplication using the heavily-armed opportunistic pathogen, *Pseudomonas aeruginosa* as a model species. We first use genomics to ask whether carrying multiple similar bacteriocins (S pyocins) increases the probability of inhibiting other strains. However, simply carrying a single uncommon S pyocin is typically as effective as carrying several. We then combine genome-editing with competition experiments to study the effects of S pyocins alone and in combinations. S pyocins sometimes function better together than alone, which helps to explain their co-carriage. However, we also find evidence of specialisation: pyocin S5 is much more effective than S2 in iron limited conditions, and vice-versa when iron is available. We test the generality of our findings using a second class of weapon carried by *P. aeruginosa*, contact-dependent inhibition, which again reveals differing effectiveness as conditions change. Our work shows how similar weapons are specialised for combat under different environmental conditions. We argue that functional specialisation is key to understanding why bacteria carry so many weapons.

## Introduction

Bacteria have evolved an immense diversity of molecular weapons they use to attack and kill competing strains and species^1–3^. Examples include many clinical antibiotics^4– 8^, but also protein bacteriocins and phage-tail derived tailocins^9^, all of which diffuse through the environment to harm competitors at a distance. At short range, bacteria attack using contact weapons such as the type VI secretion system (T6SS)^10^ and contact-dependent inhibition (CDI)^11^. Moreover, a single bacterial strain will commonly carry multiple versions of these weapons^12^. What led to bacteria evolving to be so heavily-armed? In some cases different weapons can perform different functions for bacteria: short-range weapons are particularly effective for invading a niche whereas long-range weapons may serve best to defend an established niche^13^. However, even within each category of bacterial weapon, it is common for bacteria to carry multiple similar weapons which appear to perform identical functions during combat. Why then do bacteria commonly carry such similar weapons?

Here we investigate this question using the opportunistic pathogen *Pseudomonas aeruginosa* which carries multiples of several weapon types. The type strain PAO1 produces two CDI systems, two T6SSs that deploy multiple anti-bacterial effectors, three protein bacteriocins (S pyocins), two tailocins (R/F pyocins), and a range of toxic small molecules including pyocyanin and cyanide^12^. *P. aeruginosa* is an extreme example of carrying multiple similar weapons, but the same phenomenon is seen in many other bacteria, including *E. coli*’s bacteriocins^14,15^. Here we focus on the S pyocins of *P. aeruginosa* as a model to understand the carriage of multiple similar weapons by bacteria^12^. These protein bacteriocins are released by self-lysis and use siderophore receptors to enter target cells of the same species where they kill using a toxin domain. The three S pyocins of PAO1 differ in their toxin domains, but have some overlap in their receptor-binding domains: S2 (DNAse) and S4 (tRNAse), both bind the pyoverdine receptor FpvA while S5 (pore-forming) uses the pyochelin receptor FptA^16,17^. Moreover, they are co-regulated and expressed in response to DNA damage^18^, which means they are deployed simultaneously, suggesting limited functional differentiation.

Given the lack of any apparent specialisation, a candidate explanation for carrying multiple similar weapons is to increase the probability that the carrier can kill more different potential competitors. Like many bacteriocins, immunity to S pyocins is mediated by an adjacently encoded cognate immunity protein^12^. Attacking a competing strain that carries the same S pyocin will therefore be ineffective. The potential for this and other resistance mechanisms (such as not encoding the receptor protein hijacked by the bacteriocin) may generate natural selection to carry multiple S pyocins and increase the chances that one of them is effective. Here we examine this idea using genomics but find that the effect of carrying multiple S pyocins is limited. Instead, we show that the key to targeting many strains is to carry a rare bacteriocin. We then turn to experimental work to test alternative explanations. We find that using a combination of S pyocins sometimes allows a strain to inhibit competitors more effectively than using only one. In addition, despite their similarity, S pyocins show functional specialisation, a pattern we then show is repeated in CDI systems. We conclude that, even for very similar weapons, bacteria carry multiple versions to perform distinct functions and enable combat across diverse environmental conditions.

## Results

### Carriage of multiple similar S pyocins is common in *P. aeruginosa*

Our goal is to understand why bacteria commonly carry multiple genes which encode very similar weapons. *P. aeruginosa* is an ideal model species for this question because it carries multiple similar versions of several distinct weapon types. These weapons include the S pyocins, which are protein bacteriocins homologous to the colicins of *E. coli*. The type strain *P. aeruginosa* PAO1 carries three S pyocins, S2, S4, and S5. To confirm that carriage of multiple S pyocins is common throughout *P. aeruginosa*, we systematically analysed a dataset of 517 complete *P. aeruginosa* genomes for the presence of 16 different S pyocins (Fig. 1A, S1A). This analysis reveals that S pyocins are widely distributed within the species with >90% of strains encoding at least one S pyocin. Moreover, 52% of the *P. aeruginosa* strains in the dataset encode more than one S pyocin (Fig. 1B), with up to four S pyocins found within a single strain. However, when trying to examine traits across a species, oversampling of common strains can introduce biases. To overcome this, we clustered our dataset into phylogenetically related clusters using PopPunk^19^, and treated each cluster as an independent sample. Oversampled strains will therefore be clustered into a single sample. Reassuringly, this population clustering did not greatly affect our results: 39% of the *P. aeruginosa* clusters encode more than one S pyocin (Fig. 1B, S1B).

**Figure 1:**
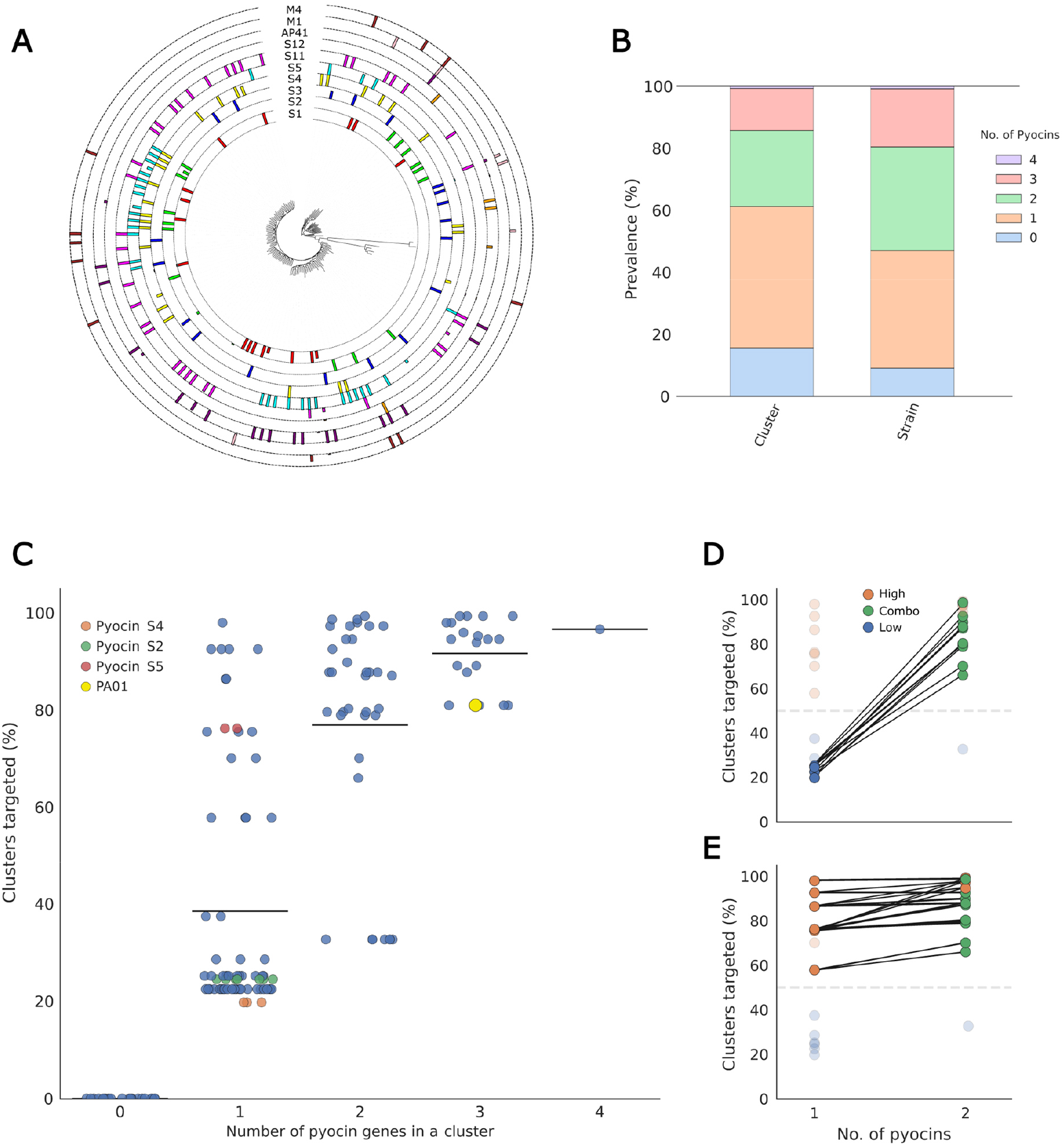
Distribution and carriage of S pyocin genes in *Pseudomonas aeruginosa* strains. (A) Whole genome phylogenetic tree and carriage of the top 10 most common S pyocin genes. (B) Proportion of strains carrying 0 - 4 pyocin genes, by PopPunk cluster and strain. (C) Number of pyocin genes vs percentage of competitors targetable (by PopPunk cluster). PAO1 is marked, and its individual S pyocins. (D) Change in percentage of competitors targetable when clusters with Low (<50% targetable) pyocins gain a second pyocin. (E) Change in percentage of competitors targetable when High (>50% targetable) pyocins gain a second pyocin.

### Improved targeting is explained by carriage of rare S pyocins

S pyocin producing cells are protected from intoxication by production of a cognate immunity protein which is encoded in the genome downstream of the toxin. In addition, non-producing cells can be protected from an attacker’s toxin by encoding so-called ‘orphan’ immunity proteins, which provide protection against a specific S pyocin without encoding the toxin. Protection can also be achieved if a cell lacks the outer membrane receptor that the S pyocin hijacks for entry into the cell. One potential reason for using multiple S pyocins against competitors, therefore, is to decrease the likelihood that the target is immune to an attack compared to attacking with just a single S pyocin.

In order to explore this hypothesis, we tested our strains of *P. aeruginosa* for i) the presence of S pyocins, ii) orphan immunity proteins, and iii) (known) outer membrane receptors targeted by S pyocins. We then used the presence and absence of these genes to predict the ability of each strain to successfully target all of the other strains in our dataset. Where the outer membrane receptor is unknown, we assumed its presence and calculated resistance based on immunity protein presence alone. We found that the ability to kill more strains (coverage) increased with the addition of more pyocins within a strain (MCMCglmm µ = 30.44, 95% CI = [28.262, 32.67], pMCMC = <0.0001]) and this trend also occurred using our population clusters (glm -(β = 28.23, SE = 0.99, t = 28.9, p = <0.0001).

This trend could be caused by additive interactions between S pyocins, with each pyocin increasing the range of strains covered, or by the increasing probability of encoding a rare pyocin where protection from immunity proteins in the population is rare. To test this hypothesis, we split our pyocins into ‘high’ (pyocins predicted to target >50% of population clusters) and ‘low’ (<50%) killing pyocins, and examined the increase in coverage provided by a second pyocin. While adding a second pyocin to a low killing pyocin does greatly increase the range of strains that could be effectively attacked, the same is not true for the high killing pyocins. Here, carriage of one of these pyocins is sufficient to be able to target the majority of other strains and carrying a second one provides little or no additional benefit (Fig. 1D).

We also assessed the hypothesis that strains may be under natural selection to carry more than one pyocin because this will make them immune to attacks from a wider range of strains (Fig. S2). This analysis suggests a modest increase in protection from attacks due to carriage of additional pyocins, which might contribute to natural selection for carriage. However, we do not expect this effect to fully explain carriage of multiple pyocins, both because it is a weak effect but also because, if this was the primary driver, we would expect strains to only carry orphan immunity loci, not the complete pair for each pyocin.

### S pyocins rarely kill more effectively together than alone

While carrying multiple S pyocins can increase the number of strains that an attacker can target, this effect is principally explained by the carriage of one low-frequency S pyocin that itself can target many strains. The carriage of additional S pyocins, particularly the more common ones, requires a different explanation. An alternative explanation is that the killing effects of pyocins combine synergistically, such that they are much more effective weapons in combination than alone. Testing this hypothesis using bioinformatics is not possible, so next we turn to experimental work using the type strain *P. aeruginosa* PAO1, which carries three S pyocins, S2, S4, and S5, to examine how they interact functionally during bacterial warfare.

We generated strains that are susceptible to each S pyocin alone and all combinations by genome editing to delete S pyocin genes and their corresponding immunity protein sequences. We then competed the wildtype (original) strain with the engineered strains to study the effects of one, two or all three S pyocins during bacterial combat in colony biofilm competitions on LB agar. This assay captures the dense, spatially-structured conditions that are typical of many bacterial communities and where bacterial weapons are thought to be most important for fitness^20^. The strains were tagged with antibiotic resistance and fluorescent protein genes to enable imaging and enumeration of the different genotypes in the competitions.

Fluorescence microscopy imaging of the colonies reveals large differences in effectiveness of the S pyocins (Fig. 2A). This is confirmed by plating and colony counts to calculate the competitive advantage provided by each pyocin alone and in combination (Fig. 2B). When starting from equal proportions (1:1), Pyocin S2 provides a ∼500-fold advantage to the attacker, while pyocins S5 (∼10-fold), and S4 (∼2-fold) provide far less of an advantage. Combinations perform similarly to the best pyocin in a pair: pyocins S2+S4 and S2+S5 do not provide a significantly different advantage compared to pyocin S2 by itself. Similarly, S4+S5 does not provide a significantly different advantage to S5 alone. Finally, the combination of all three was not significantly different from either combination containing S2, or S2 by itself. We also tested competition outcomes when the attacker had a frequency advantage (initial ratio of 10:1, Fig. S5). Here, competitive advantages are generally higher and there is one combination (S2+S5) that is significantly better together than alone. These trends were the same at the edge of the colony (Fig. S5). There is some evidence, therefore, that S pyocins can kill more effectively together than alone but this occurs in a minority of cases studied. This suggests that additional explanations are needed to explain why *P. aeruginosa* carries multiple S pyocins.

**Figure 2:**
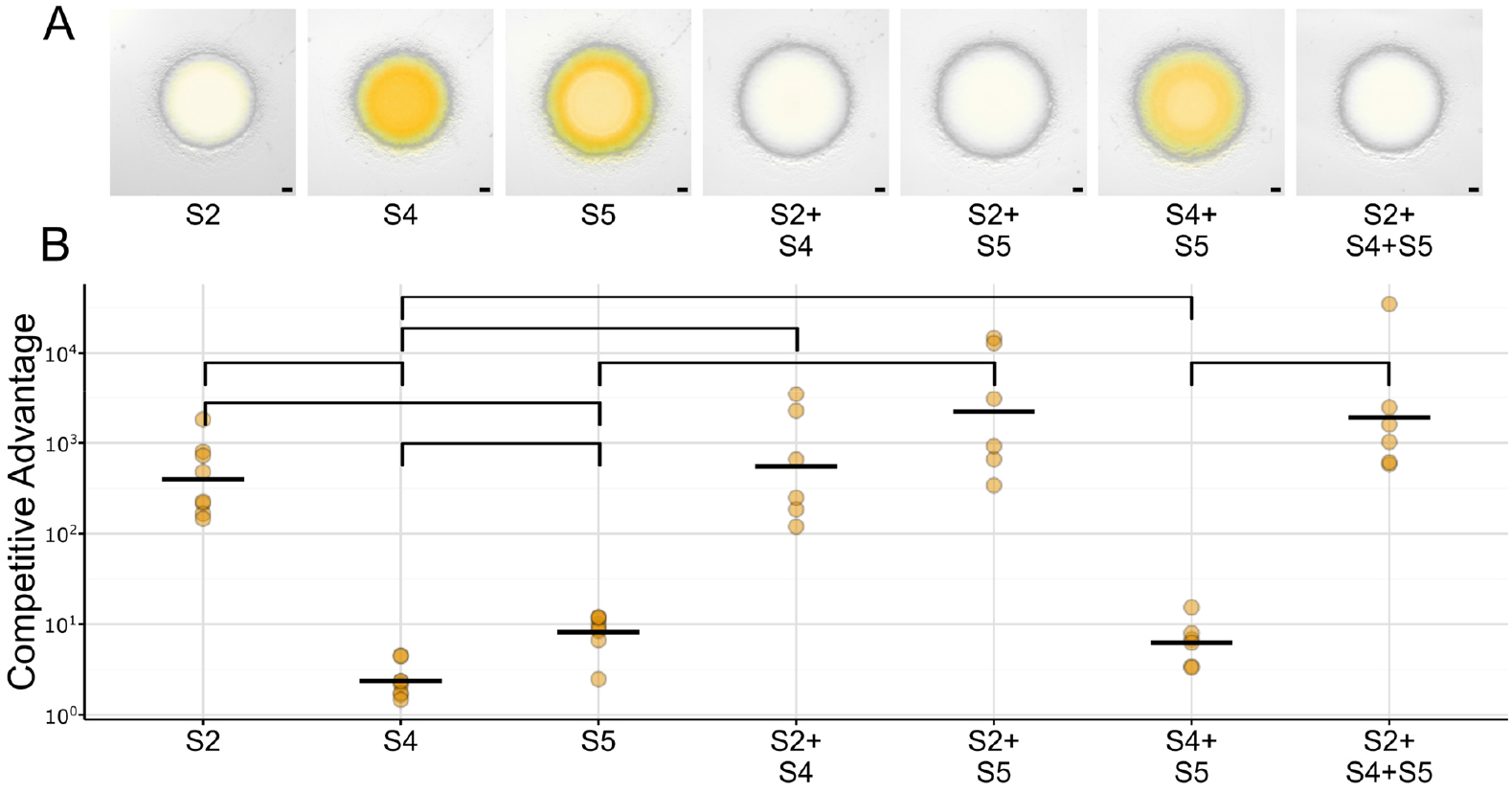
Competitive advantage provided by single and combinations of the S pyocins of *P. aeruginosa* PAO1 in colony biofilms. (A) Microscopy images of colony competitions carried out between equal proportions (1:1) of wild-type (unlabelled) and fluorescently-tagged S pyocin susceptible mutants (orange). Mutant strains had either single or combinations of deletions of both the toxin and immunity protein of pyocins S2, S4, and/or S5, and were tagged with Tn7-GmR-mScarlet/eYFP. Wild-type was untagged, images show a merger of the brightfield and mutant strain fluorescence (orange). (B) Corresponding competitive advantage determined by spot-plating and counting numbers of CFU. Brackets indicate significant differences (p < 0.05) according to BH-corrected pairwise t-tests.

### Different S pyocins function best under different conditions

A striking pattern in our experimental data is that two of the pyocins (S4 and S5) perform very poorly relative to the other (pyocin S2) in competition experiments. This pattern raises the interesting possibility that the other pyocins have not evolved to function well under the conditions of our experiment. Pyocins S2 and S4 gain entry into target cells via the pyoverdine transporter FpvA, while S5 enters through the pyochelin transporter FptA. Production of these siderophores and their receptors varies depending on iron availability^21,22^. We, therefore, decided to test the effectiveness of each pyocin in media supplemented with the iron chelator bipyridyl to create an environment where iron is limited. Using the colony biofilm competition assay, these experiments reveal that pyocin S2 becomes much less effective when iron is limited, providing only a ∼10-fold advantage. By contrast, pyocin S5 becomes much more effective, now providing a ∼100-fold advantage (Fig. 3AB). Thus, an attacker using both pyocins S2 and S5 has the potential to strongly suppress a competitor in environments where the levels of iron differ. Under iron-limited conditions, we also see a number of additional cases where two pyocins in combination can be more effective at suppressing a target strain than either one alone e.g. S2+S5 gives a better competitive advantage than either S2 or S5 alone (Fig. S5). These data indicate that both improved killing and functional specialisation provide benefits to carrying multiple similar bacteriocins.

**Figure 3:**
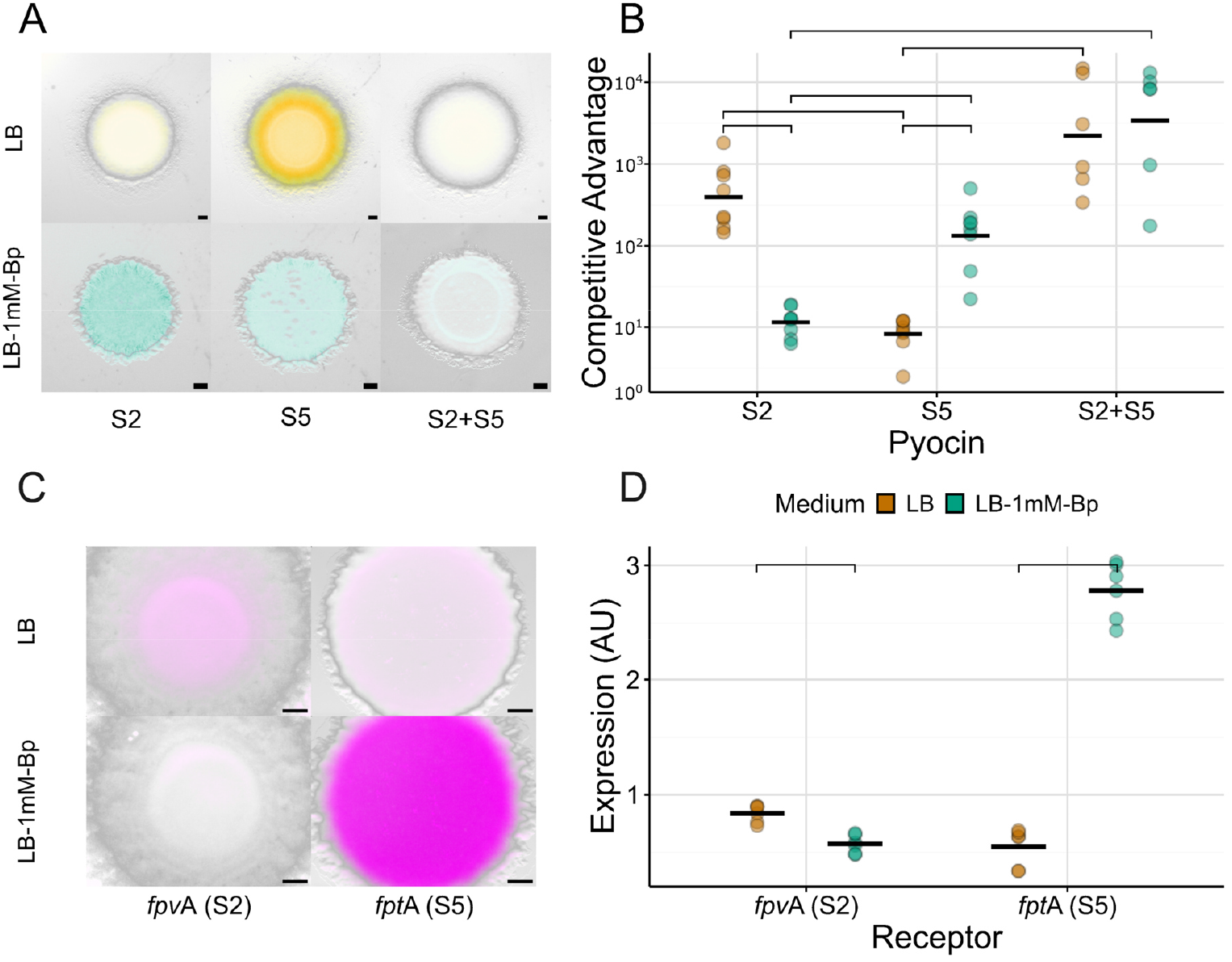
Competitive advantage provided by pyocins S2 and/or S5 and expression of corresponding receptors FpvA and FptA on LB or LB 1mM Bipyridyl. (A) Microscopy images of colony competitions carried out between equal proportions (1:1) of wild-type and S pyocin susceptible mutants of *P. aeruginosa* PAO1. Mutant strains had either single or combinations of deletions of both the toxin and immunity protein of pyocins S2 and/or S5, and were tagged with Tn7-GmR-mScarlet/eYFP. Wild-type was untagged, images show a merger of the brightfield and mutant strain fluorescence on either LB 1.5% agar (orange, top row) or LB 1.5% agar + 1mM bipyridyl (Bp) (blue, bottom row). (B) Corresponding competitive advantage determined by spot-plating and counting numbers of CFU. (C) Microscopy images of colonies of *P. aeruginosa* PAO1 carrying plasmids showing expression of FpvA or FptA. Strains were transformed with plasmids pOPC-244 or pOPC-248^23^, then grown on either LB 1.5% agar or LB 1.5% agar + 1mM bipyridyl. Signal from YPet (magenta) corresponds to expression of the siderophore receptor which is hijacked for pyocin import. (D) Quantification of siderophore receptor expression microscopy images. Receptor-YPet signal was normalized to *pilM*-mCherry signal. Brackets indicate significant differences (p < 0.05) according to BH-corrected pairwise t-tests.

To enter a cell, S pyocins must bind to and pass through a siderophore transporter protein on the target cell. One candidate explanation for changes in their effectiveness across conditions, therefore, is that the target receptors are expressed differently across conditions. To explore this hypothesis, we quantified the expression of the two different siderophore transporter proteins using fluorescent reporters^23^, which fuse the promoters of *fpvAI* or *fptA* to mCherry. Imaging of colonies grown under the same conditions as our competition assays suggests that there is indeed differential expression of the two transporters across the two conditions (Fig. 3C). We, therefore, quantified the mCherry signal and normalized it against constitutively expressed YPet to account for any variations in growth. This analysis shows that *fpvAI* (pyocin S2 receptor) expression is highest without iron limitation while *fptA* (pyocin S5 receptor) expression is greatly increased and higher under iron limitation (Fig. 3D). Overall, these data suggest that the different pyocins perform best under different conditions, and that these differences are explained by differential expression of the receptors in target strains.

### Similar patterns are seen for contact-dependent inhibition

Our findings with S pyocins support the importance of specialisation of weapon function, where it is beneficial for a cell to carry multiple versions of a given weapon to help ensure it has the right weapon for a given condition. To test the generality of this finding, we turned to a second type of bacterial weapon: contact-dependent inhibition (CDI). *P. aeruginosa* PAO1 carries two similar but distinct CDI systems, CDI1 and CDI2^24^. Starting with genomics again, our analysis shows that most strains have CDI1 (Fig. 4A), and almost 50% of strains also have CDI2 (Fig. 4B) where carriage of the second system is distributed widely across the species. Predicting attacker success from sequence data is more challenging for CDI systems than for S pyocins. Firstly, immunity genes are harder to identify and assign because frequent recombination in the cytotoxic domains degrades the regions with the immunity genes. Secondly, the outer membrane receptors on target cells that are used by CDI1 and CDI2 to import the toxin are unknown (they are likely encoded by essential genes because they have not been identified through transposon screens^25^). As a result, our predictions here on whether a strain can effectively intoxicate another strain is based solely on possession of cytotoxic domains (immunity genes and receptors are not used as they were for the S pyocins). However, there is a large variety of different cytotoxic domains associated with these two systems (38) compared to S pyocin (12) toxin domains (Fig. 4C), which helps to improve the resolution of predictions. Similarly to carrying a rare S pyocin, possession of one weapon appears to be enough to target over 80% of competitors whereas there was only a slight increase in targeting breadth through the addition of a second CDI system (Fig. 4D). Carrying multiple similar CDI systems, therefore, does not appear to be explained by the need to increase targeting range.

**Figure 4:**
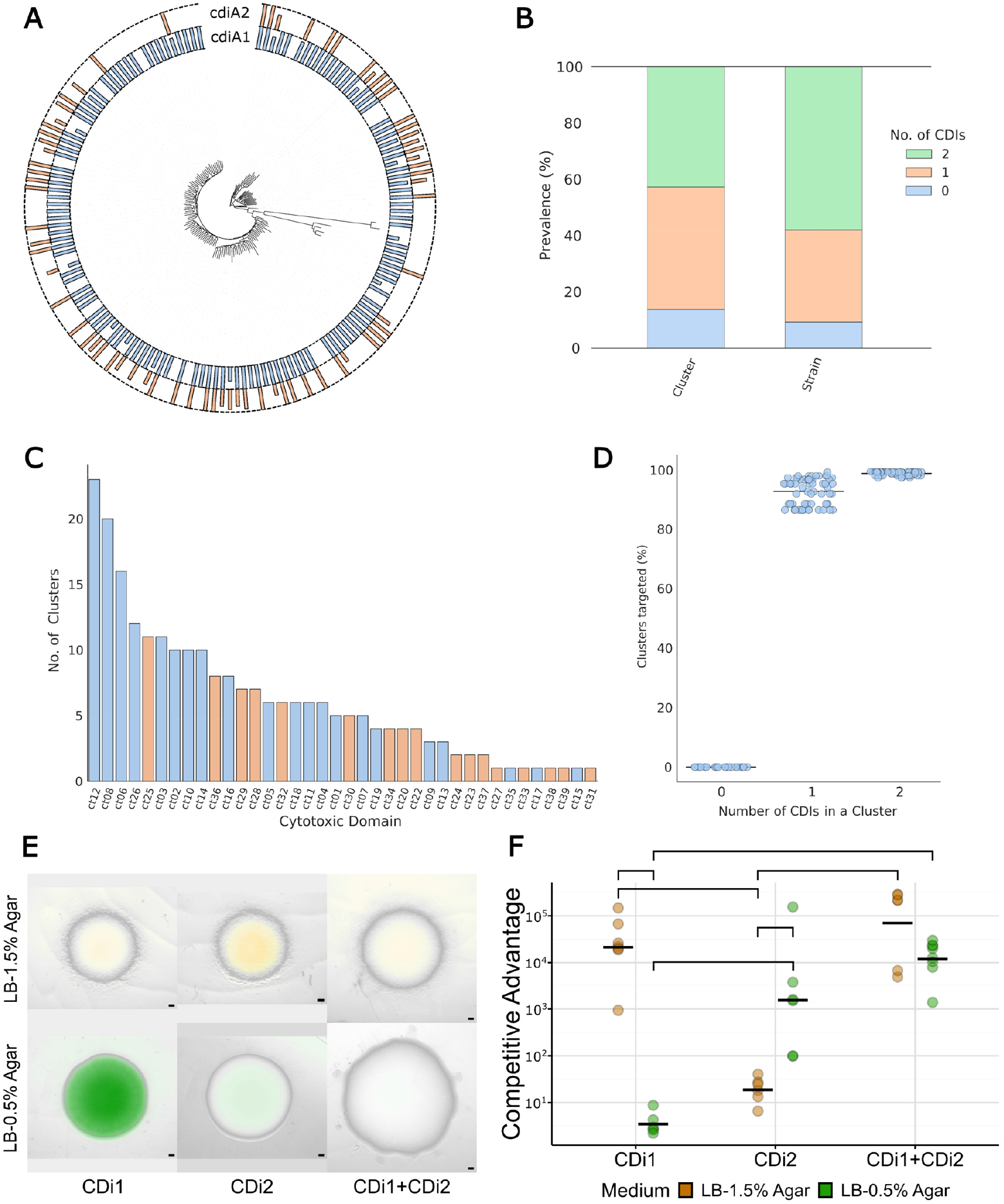
Distribution and carriage of CDI genes in Pseudomonas aeruginosa strains. (A) Whole genome phylogenetic tree and carriage *cdi*A1 and *cdi*A2. (B) Proportion of strains carrying 0 - 2 *cdiA* genes, by PopPunk cluster and strain. (C) Number of PopPunk clusters carrying different cdiA cytotoxic domains. (D) Number of CDI genes vs percentage of competitors targetable. (E) Microscopy images of colony competitions carried out between wild-type and S pyocin susceptible mutants of *P. aeruginosa* PAO1. Mutant strains had either single or combinations of deletions of both the CDI filament and immunity protein of CDI1, and/or CDI2, and were tagged with Tn7-GmR-mScarlet/eYFP. Wild-type was untagged, images show a merger of the brightfield and mutant strain fluorescence (1.5% agar, orange; 0.5% agar, green). (F) Corresponding competitive advantage determined by spot-plating and counting numbers of CFU. Brackets indicate significant differences (p < 0.05) according to BH-corrected pairwise t-tests.

We, therefore, again moved to experiments using single and double deletion mutants of *P. aeruginosa* PAO1 susceptible to one or both CDI systems. Previous work shows that effectiveness and expression of contact dependent weapons depends strongly on the physical environment, for example, shaking vs static liquid culture influences CDI expression^24^ and contact weapons are generally more effective on surfaces than in liquid culture^26^. To explore conditions that might change the impacts of PAO1’s two CDI systems without undermining their effectiveness, we decided to assess the effectiveness of CDI 1 and 2 in colony biofilms grown on hard (1.5%) and soft (0.5%) agar (Fig. 4E). Consistent with the importance of the physical environment, CDI1 provided a 10,000-fold advantage on hard agar, but no advantage at all on soft agar (Fig. 4F). Conversely, CDI2 provided a 10-fold advantage on hard agar which increased to 1,000-fold on soft agar. We again observe conditional specialisation, therefore, and also combinatorial effectiveness in some situations (Fig. S6), which mirrors our findings for the S pyocins.

## Discussion

Bacteria often carry multiple weapons for attacking competitors. While some weapons appear to be distinct and functionally specialised, there are many examples where bacteria carry multiple versions of weapons that are very similar. Here we show that despite their similarity, these seemingly redundant weapons can also perform distinct functions. Focussing first on the bacteriocins of *P. aeruginosa*, we show that it is common for a strain to carry multiple S pyocins, which can differ greatly in their effectiveness as a function of iron availability. Carrying more than one pyocin, therefore, allows a strain to effectively attack competitors across a wider range of environmental conditions than carrying only one. In addition, in some conditions, two pyocins used in combination are more effective than one at suppressing the competing strain, giving a second advantage of co-carriage. We observed the same patterns for contact dependent weapon (CDI), where agar concentration differentiated the performance of the two CDI systems and a strain wielding both gained a large advantage under both conditions.

Both S pyocins and CDI rely on cell-surface proteins to gain access to their cellular targets^27^. The expression of outer membrane proteins is regulated by multiple environmental factors^28^, which means the abundance of the surface proteins used as a receptor by a specific weapon is often tightly linked to environmental conditions. While S pyocins bind their receptors with high affinity^29^ and likely the same for CDI^30^, a weapon can become useless under conditions where the receptor is not expressed. By carrying multiple weapons that target different receptors, bacteria can thus ensure that they can attack their competitors under different environmental conditions. In support of this effect, we observed that expression of the pyochelin receptor, FptA, was greatly increased under low-iron conditions, and that under these conditions pyocin S5 (which hijacks this receptor) provided a large competitive advantage. Though the outer membrane receptors for PAO1’s CDI1 and CDI2 are unknown, agar concentration could be influencing the expression of outer-membrane proteins, similar to the osmolarity-sensitive OmpC/OmpF^31^ which are used as CDI receptors in *E. coli*^32^.

Bacteria use the Type Six Secretion System to simultaneously inject multiple toxin effectors directly into competitor cells^10^. There are differences in this example to the carriage of multiple similar weapons because the distinct toxin effectors are all delivered together by a shared apparatus that physically punctures the target cell. Co-carriage of effectors, therefore, is not explained by the need to always have a toxin for which there are receptors on the target cell. However, there is evidence that some effectors function better than others based on environmental conditions^33^ and there is evidence that effectors can be better at killing in combination than alone, which are both consistent with our findings. Similar patterns are observed for pathogenic bacteria, which carry large numbers of different effector proteins that are directly injected into eukaryotic cells. When *Legionella pneumophila* infects amoeba/macrophage^34^ and *P. syringae* infects plant cells^35,36^, removal of a single effector often has no effect, suggesting that that these effectors have overlapping, redundant functions. However, careful experimentation has revealed that these effectors are not truly redundant and are more likely adapted for optimal use in different hosts. These examples suggest that our observations are part of a broader pattern where functional specialisation and combinational effects lead bacteria to carry multiple versions of a particular protein or molecular system.

Bacteria have evolved a wide variety of different weaponry, and many strains carry large, seemingly redundant arsenals^1^. Our work suggests that these weapons are not redundant but instead have evolved for specialized use under specific conditions, enabling effective combat under a wide variety of environmental conditions.

## Methods

*P. aeruginosa* genomes were accessed from the BV-BRC database^37^. Complete genomes were selected with ‘good’ genome quality. Genomes were removed if they were less than 5 Mbp in length or had a low CheckM completeness score. Genomes were also removed by investigating the relationship between genome length and number of coding sequences. For any one species, this relationship is linear, therefore, genomes which deviated from this linear relationship by > 500 coding sequences were removed^38^. Finally, genomes with suspected contamination were removed using the panaroo-qc tool, which uses mash to identify distances between genomes and a database of refseq genomes^39,40^. This produced a database of 517 *P. aeruginosa* genomes.

### *P. aeruginosa* phylogeny

The pangenome of 517 *P. aeruginosa* genomes was calculated using panaroo with a core gene threshold of 99% and ‘sensitive’ mode to avoid removing rare pyocins. The 4558 core genes were aligned using MAFFT^41^. The phylogeny was calculated using iqtree2 with the general time reversible model (GTR+F+R10)^42^. The resulting phylogenetic tree was visualised using iTOL^43^. To assess the genetic diversity and population structure of our dataset, genomes were clustered using popPunk with the --max-zero-dist flag and the *P. aeruginosa* popPunk dataset v1^44^.

### Identifying pyocin receptors

To identify strains encoding different pyocin outer membrane receptors, a database of known receptors was built (Table S1) and all genomes scanned using abricate (https://github.com/tseemann/abricate) with default coverage and identity thresholds.

### Identifying pyocins

To identify pyocins, open reading frames were predicted using getORF from EMBOSS^45^. Pfam profiles of pyocin cytotoxic domains and their cognate immunity proteins were used to identify potential pyocins using HMMer3.0^46^ (Table S1). Immunity proteins for M1 and M4 could not be detected using Pfam domains, but were identified using Blastp to identify pmiC from *P. aeruginosa* BL03 (Q057_04090) and *pmiA* from *P. aeruginosa* F291007 (K5B02_RS22775).

Pyocin immunity proteins can share similar domains with many toxin/antitoxin systems and they can be difficult to identify without the context of a cognate toxin. To identify orphan immunity proteins, we made a database of all immunity protein genes found downstream of a cognate toxin. We then used abricate to scan our dataset for new immunity proteins without a cognate toxin.

### Identifying CDI systems

To identify CDI toxins, all open reading frames were scanned for Pfam domains associated with CDI toxins (cdiA1: PA0041 and cdiA2: PA2642) and transporters (cdiB1: PA0040 and cdiB2: PA2463) from *P. aeruginosa* PA01 (Table S1). Possible CDI systems contained a protein with all cdiA profiles associated with a predicted transporter with all cdiB profiles. Possible cdiA genes were then scanned for 5 known *P. aeruginosa* cdiA receptor-binding domains^47^ using BLAST.

To identify the different toxins used by CDI systems, we scanned our predicted cdiA genes for 39 known CDI toxin domains in *P. aeuginosa*^47^ using BLASTp. We filtered and removed hits which were less than 10% of the maximum score observed for that cytotoxic domain and used the best cytotoxic domain prediction for each cdiA gene.

### Predicting weapon spectrum

#### Pyocins

For a pyocin to kill a competitor, that competitor needs to express the correct receptor and not encode the immunity protein for that pyocin. A combination of pyocins, receptors and orphan immunity proteins was used to predict whether each strain could target every other strain. For pyocins with no known receptor, we assumed that the receptor is found within all strains. For pyocin S12, a family of orphan immunities that are not annotated as reading frames and have been shown to fail to provide protection was removed from analysis^48^. To make predictions for each popPUNK cluster, the most common combination of pyocins, immunity and receptors was identified for each cluster and used to represent the cluster (Fig. S3).

#### CDI

CDI systems may contain many degraded cytotoxic domains and immunity genes downstream of the toxin. As it can be difficult to determine if they are expressed and could provide protection, CDI killing spectrum was determined solely by the combination of 39 cytotoxic domains in the cdiA1/2 toxins. As many CDI receptors are unknown, they were also not used to account for CDI targeting spectrum. To make predictions for each popPUNK cluster, the most common combination of CDI cytotoxic domains was identified for each cluster and used to represent the cluster (Fig. S4).

### Experiments

#### Strain construction

Strains used in this study are summarized in Table S2. Deletion mutants were constructed as previously using two-step allelic exchange with pEXG2 and conjugating with *E. coli* JKE201^49–51^. Pyocin susceptible strains were constructed in two steps, first the pyocin was deleted, then the immunity to prevent the susceptible strain from being immediately killed during the construction process. The double and triple mutants were made by making sequential deletions. Primer sequences are listed in Table S3. Mutants were confirmed by Sanger sequencing (Source Bioscience, Nottingham, UK). Strains were subsequently tagged with constitutively expressed eYFP and mScarlet using pUC18-mini-Tn7-GmR^52^. Equal numbers of replicates were performed with each attacker/susceptible combination carrying each fluorescent marker. Strains are available upon request.

#### Culturing

Strains were recovered from freezer stocks by streaking onto LB 1.5% agar, then incubating overnight at 30°C. Cells were scraped off the overnight plate and resuspended to an initial OD_600_ of 1.0, then mixed at defined ratios, and 1 µL spotted on media. For the CDI competitions, inocula were serially diluted 10-fold three times as CDI was previously found to be more effective at lower densities^53^. Plates of LB 1.5% agar, LB 0.5% agar, and LB 1.5% agar + 1mM bipyridyl were prepared immediately prior to use and dried for 15 minutes in a laminar flow hood. The density of the inoculum was determined by serial dilution followed by spot plating; inoculum densities of OD_600_ 1.0 were found to correspond to approximately 2x10^6^ cells per 1 μL droplet of inoculum. Data presented are from independent colonies (biological replicates). Data presented in Fig. 4 are from initial density of 10^-2^, and all three densities tested are presented in Fig. S6.

#### Quantification of competition outcomes

After 48 h of growth at room temperature, images of colonies were taken using a Zeiss Axio Zoom V16 microscope with a Zeiss MRm camera, 0.5X PlanApo Z air objective and HXP 200C fluorescence light source. Colonies of different initial density and attacker:sensitive ratios were imaged at the same magnification. To make the composite images shown in the figures, the display histograms of each channel were normalized to the set of colonies sharing the same media, meaning images can be compared within the same media but not between. After imaging, colonies were destructively sampled with a 10 µL pipette tip at the centre and edge of the colony into 0.9% saline. The centre includes the initial inoculation zone, while the edge was sampled in an arc shape from the exterior-most 1-2mm of the colony. Samples were homogenized, serially diluted 10-fold and 5 μL spotted onto LB and LB + 50 μg/mL gentamycin and incubated at 30°C overnight, then colonies counted.

#### Calculation of competitive advantage

Using the original density counted from the initial inoculum cultures and the known inoculum ratios, the initial ratio of attacker:susceptible strains was determined. CFU counts of the serially diluted centre and edge samples were used to determine the final ratio, with a detection limit of 10 CFU/mL used to replace zeros and prevent dividing by zero. Competitive advantage was calculated as the final ratio / initial ratio of attacker:susceptible cells and is plotted on log axes. Competitive advantages were compared using pairwise t-tests calculated using ggpubr, with Benjamini-Hochberg correction for multiple testing^54^.

#### Quantification of receptor expression

Expression of *fpvA* and *fptA* was measured using plasmids with the promoters for these genes fused to the fluorescent protein YPet, and the constitutive *pilM* promoter fused to mCherry^23^. Plasmids pOPC-244 and pOPC-248 were conjugated into wild-type PAO1 as above. Colonies were inoculated as for competition assays, but only using the single strain, and 50 μg / mL Gm was included in the freshly poured agar plates to maintain the reporter plasmids. Colonies were imaged after 48 h of growth in both red and yellow channels using a Zeiss Axio Zoom V16 microscope with a Zeiss MRm camera, 0.5X PlanApo Z air objective and HXP 200C fluorescence light source. For displaying images of the colonies, yellow fluorescence was scaled to the minimum and maximum obtained from all displayed micrographs and merged with the brightfield. For quantification, the yellow signal was normalized to the red signal. Data represent 6 biological replicates.

### Data availability

Data for this work is available at https://doi.org/10.6084/m9.figshare.31450261.

## Acknowledgements

We thank Olivier P. Cunrath for providing the promoter-fusion plasmids, and Jacob Palmer and Erik Bakkeren for feedback.

## References

1. Granato, E. T., Meiller-Legrand, T. A. & Foster, K. R. The evolution and ecology of bacterial warfare. Current Biology 29, R521–R537 (2019).

2. García-Bayona, L. & Comstock, L. E. Bacterial antagonism in host-associated microbial communities. Science 361, eaat2456 (2018).

3. Hibbing, M. E., Fuqua, C., Parsek, M. R. & Peterson, S. B. Bacterial competition: surviving and thriving in the microbial jungle. Nat Rev Microbiol 8, 15–25 (2010).

4. Clardy, J., Fischbach, M. & Currie, C. The natural history of antibiotics. Curr Biol 19, R437–R441 (2009).

5. Dulmage, H. T. The Production of Neomycin by Streptomyces fradiae in Synthetic Media. Appl Microbiol 1, 103–106 (1953).

6. Schatz, A., Bugle, E. & Waksman, S. A. Streptomycin, a Substance Exhibiting Antibiotic Activity Against Gram-Positive and Gram-Negative Bacteria. Proceedings of the Society for Experimental Biology and Medicine 55, 66–69 (1944).

7. Westhoff, S. et al. Spatial structure increases the benefits of antibiotic production in Streptomyces. Evolution 74, 179–187 (2020).

8. Wright, E. S. & Vetsigian, K. H. Inhibitory interactions promote frequent bistability among competing bacteria. Nat Commun 7, 11274 (2016).

9. Nakayama, K. et al. The R-type pyocin of Pseudomonas aeruginosa is related to P2 phage, and the F-type is related to lambda phage. Molecular Microbiology 38, 213–231 (2000).

10. Hachani, A. et al. Type VI Secretion System in Pseudomonas aeruginosa. J Biol Chem 286, 12317–12327 (2011).

11. Aoki, S. K. et al. Contact-Dependent Inhibition of Growth in Escherichia coli. Science 309, 1245–1248 (2005).

12. Ghequire, M. G. K. & De Mot, R. Ribosomally encoded antibacterial proteins and peptides from Pseudomonas. FEMS Microbiology Reviews 38, 523–568 (2014).

13. Booth, S. C., Smith, W. P. J. & Foster, K. R. The evolution of short- and long-range weapons for bacterial competition. Nat Ecol Evol 7, 2080–2091 (2023).

14. Sharp, C. & Foster, K. Bacterial warfare is associated with virulence and antimicrobial resistance. 2024.11.06.622277 Preprint at 10.1101/2024.11.06.622277 (2024).

15. Gordon, D. M. & O’Brien, C. L. Bacteriocin diversity and the frequency of multiple bacteriocin production in Escherichia coli. Microbiology 152, 3239–3244 (2006).

16. Denayer, S., Matthijs, S. & Cornelis, P. Pyocin S2 (Sa) Kills Pseudomonas aeruginosa Strains via the FpvA Type I Ferripyoverdine Receptor. Journal of Bacteriology 189, 7663–7668 (2007).

17. Elfarash, A. et al. Pore-forming pyocin S5 utilizes the FptA ferripyochelin receptor to kill Pseudomonas aeruginosa. Microbiology (Reading) 160, 261–269 (2014).

18. Matsui, H., Sano, Y., Ishihara, H. & Shinomiya, T. Regulation of pyocin genes in Pseudomonas aeruginosa by positive (prtN) and negative (prtR) regulatory genes. J Bacteriol 175, 1257–1263 (1993).

19. Lees, J. A. et al. Fast and flexible bacterial genomic epidemiology with PopPUNK. Genome Res 29, 304–316 (2019).

20. Flemming, H.-C. & Wuertz, S. Bacteria and archaea on Earth and their abundance in biofilms. Nat Rev Microbiol 17, 247–260 (2019).

21. Cunrath, O. et al. A cell biological view of the siderophore pyochelin iron uptake pathway in seudomonas aeruginosa. Environmental Microbiology 17, 171–185 (2015).

22. Dumas, Z., Ross-Gillespie, A. & Kümmerli, R. Switching between apparently redundant iron-uptake mechanisms benefits bacteria in changeable environments. Proc Biol Sci 280, 20131055 (2013).

23. Ferry, M. et al. Characterisation of Pseudomonas aeruginosa’s metal-responsive TonB-dependent transporters. 2024.10.14.618150 Preprint at 10.1101/2024.10.14.618150 (2024).

24. Mercy, C., Ize, B., Salcedo, S. P., Bentzmann, S. de & Bigot, S. Functional characterization of Pseudomonas contact dependent growth inhibition (CDI) systems. PLOS ONE 11, e0147435 (2016).

25. Allen, J. P. et al. A comparative genomics approach identifies contact-dependent growth inhibition as a virulence determinant. Proceedings of the National Academy of Sciences 117, 6811–6821 (2020).

26. Wood, T. E. et al. The Pseudomonas aeruginosa T6SS Delivers a Periplasmic Toxin that Disrupts Bacterial Cell Morphology. Cell Rep 29, 187-201.e7 (2019).

27. Nolan, L. M. & Allsopp, L. P. Antimicrobial Weapons of Pseudomonas aeruginosa. in Pseudomonas aeruginosa: Biology, Pathogenesis and Control Strategies (eds Filloux, A. & Ramos, J.-L.) 223–256 (Springer International Publishing, Cham, 2022). doi:10.1007/978-3-031-08491-1_8.

28. Chevalier, S. et al. Structure, function and regulation of Pseudomonas aeruginosa porins. FEMS Microbiology Reviews 41, 698–722 (2017).

29. White, P. et al. Exploitation of an iron transporter for bacterial protein antibiotic import. Proceedings of the National Academy of Sciences 114, 12051–12056 (2017).

30. Ruhe, Z. C. et al. CdiA Effectors Use Modular Receptor-Binding Domains To Recognize Target Bacteria. mBio 8, 10.1128/mbio.00290-17 (2017).

31. Forst, S. A. & Roberts, D. L. Signal transduction by the EnvZ-OmpR phosphotransfer system in bacteria. Research in Microbiology 145, 363–373 (1994).

32. Csonka, L. N. Physiological and genetic responses of bacteria to osmotic stress. Microbiol Rev 53, 121–147 (1989).

33. LaCourse, K. D. et al. Conditional toxicity and synergy drive diversity among antibacterial effectors. Nat Microbiol 3, 440–446 (2018).

34. O’Connor, T. J., Adepoju, Y., Boyd, D. & Isberg, R. R. Minimization of the Legionella pneumophila genome reveals chromosomal regions involved in host range expansion. Proceedings of the National Academy of Sciences 108, 14733–14740 (2011).

35. Cunnac, S. et al. Genetic disassembly and combinatorial reassembly identify a minimal functional repertoire of type III effectors in Pseudomonas syringae. Proceedings of the National Academy of Sciences 108, 2975–2980 (2011).

36. Kvitko, B. H. et al. Deletions in the Repertoire of Pseudomonas syringae pv. tomato DC3000 Type III Secretion Effector Genes Reveal Functional Overlap among Effectors. PLOS Pathogens 5, e1000388 (2009).

37. Olson, R. D. et al. Introducing the Bacterial and Viral Bioinformatics Resource Center (BV-BRC): a resource combining PATRIC, IRD and ViPR. Nucleic Acids Res 51, D678–D689 (2022).

38. Horesh, G. et al. A comprehensive and high-quality collection of Escherichia coli genomes and their genes. Microbial Genomics 7, 000499 (2021).

39. Tonkin-Hill, G. et al. Producing polished prokaryotic pangenomes with the Panaroo pipeline. Genome Biol 21, 180 (2020).

40. Ondov, B. D. et al. Mash: fast genome and metagenome distance estimation using MinHash. Genome Biol 17, 132 (2016).

41. Katoh, K., Misawa, K., Kuma, K. & Miyata, T. MAFFT: a novel method for rapid multiple sequence alignment based on fast Fourier transform. Nucleic Acids Res 30, 3059–3066 (2002).

42. Minh, B. Q. et al. IQ-TREE 2: New Models and Efficient Methods for Phylogenetic Inference in the Genomic Era. Mol Biol Evol 37, 1530–1534 (2020).

43. Letunic, I. & Bork, P. Interactive Tree Of Life (iTOL) v5: an online tool for phylogenetic tree display and annotation. Nucleic Acids Res 49, W293–W296 (2021).

44. Blackwell, G. A. et al. Exploring bacterial diversity via a curated and searchable snapshot of archived DNA sequences. PLOS Biology 19, e3001421 (2021).

45. Rice, P., Longden, I. & Bleasby, A. EMBOSS: The European Molecular Biology Open Software Suite. Trends in Genetics 16, 276–277 (2000).

46. Eddy, S. R. Accelerated Profile HMM Searches. PLOS Computational Biology 7, e1002195 (2011).

47. Allen, J. P. & Hauser, A. R. Diversity of contact-dependent growth inhibition systems of Pseudomonas aeruginosa. J Bacteriol 201, e00776–18 (2019).

48. McCaughey, L. C. et al. Discovery, characterization and in vivo activity of pyocin SD2, a protein antibiotic from Pseudomonas aeruginosa. Biochem J 473, 2345–2358 (2016).

49. Choi, K.-H. & Schweizer, H. P. An improved method for rapid generation of unmarked Pseudomonas aeruginosa deletion mutants. BMC Microbiology 5, 30 (2005).

50. Rietsch, A., Vallet-Gely, I., Dove, S. L. & Mekalanos, J. J. ExsE, a secreted regulator of type III secretion genes in Pseudomonas aeruginosa. Proc Natl Acad Sci U S A 102, 8006–8011 (2005).

51. Harms, A. et al. A bacterial toxin-antitoxin module is the origin of inter-bacterial and inter-kingdom effectors of Bartonella. PLoS Genet 13, e1007077 (2017).

52. Choi, K.-H. & Schweizer, H. P. mini-Tn7 insertion in bacteria with single attTn7 sites: example Pseudomonas aeruginosa. Nat Protoc 1, 153–161 (2006).

53. Booth, S. C., Meacock, O. J. & Foster, K. R. Cell motility empowers bacterial contact weapons. The ISME Journal 18, wrae141 (2024).

54. Alboukadel Kassambara. ggpubr: ‘ggplot2’ Based Publication Ready Plots. (2025).

