## SupplementalFigures for "Similar weapons play different roles in bacterial competition"

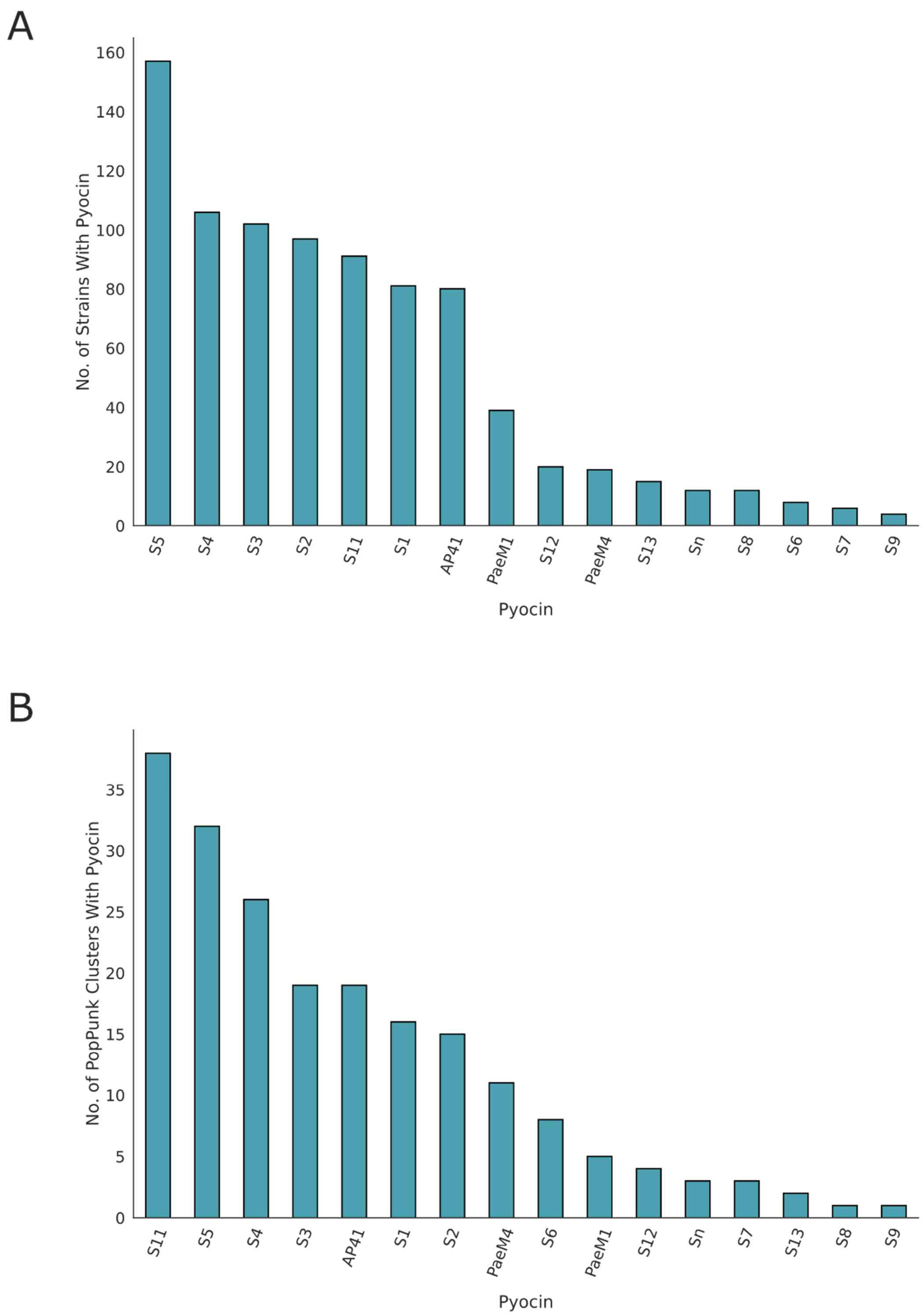

Supplementary Figure 1: Distribution of pyocins in 517 strains of *P. aeruginosa*. A) Number of strains containing each pyocins. B) Number of popPunk clusters with each pyocin.

A

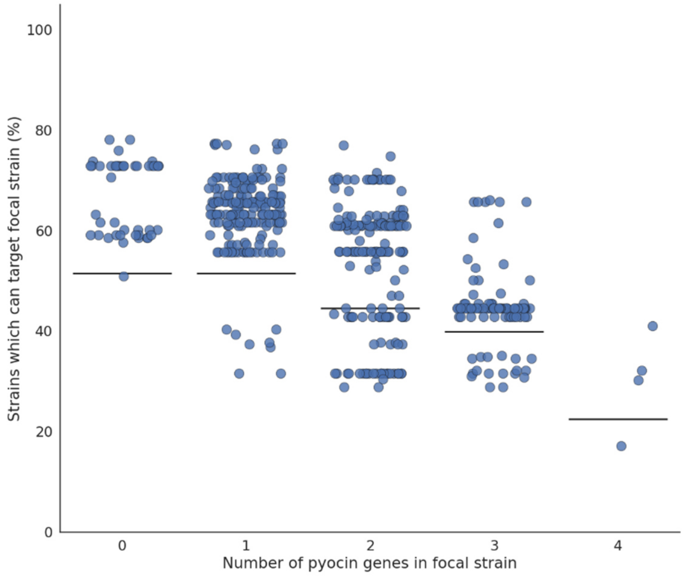

B

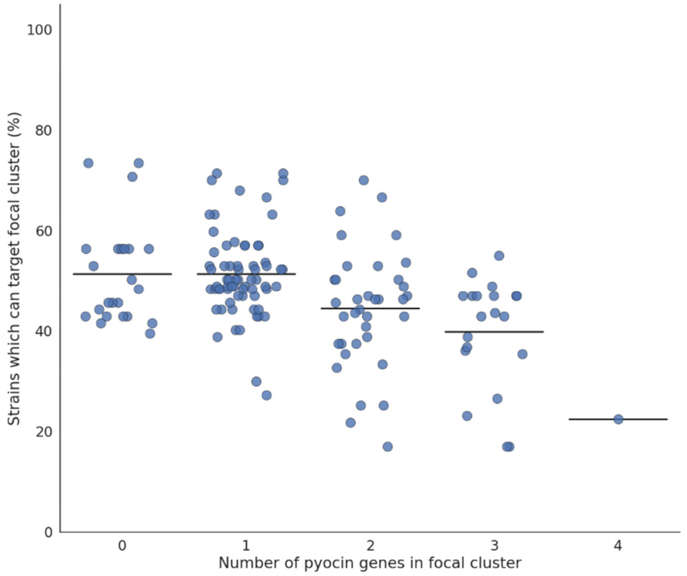

Supplementary Figure 2: Number of pyocin genes and the percentage of competitors with can target that strain/cluster. Data are shown by Strain (A) and by PopPunk cluster (B).

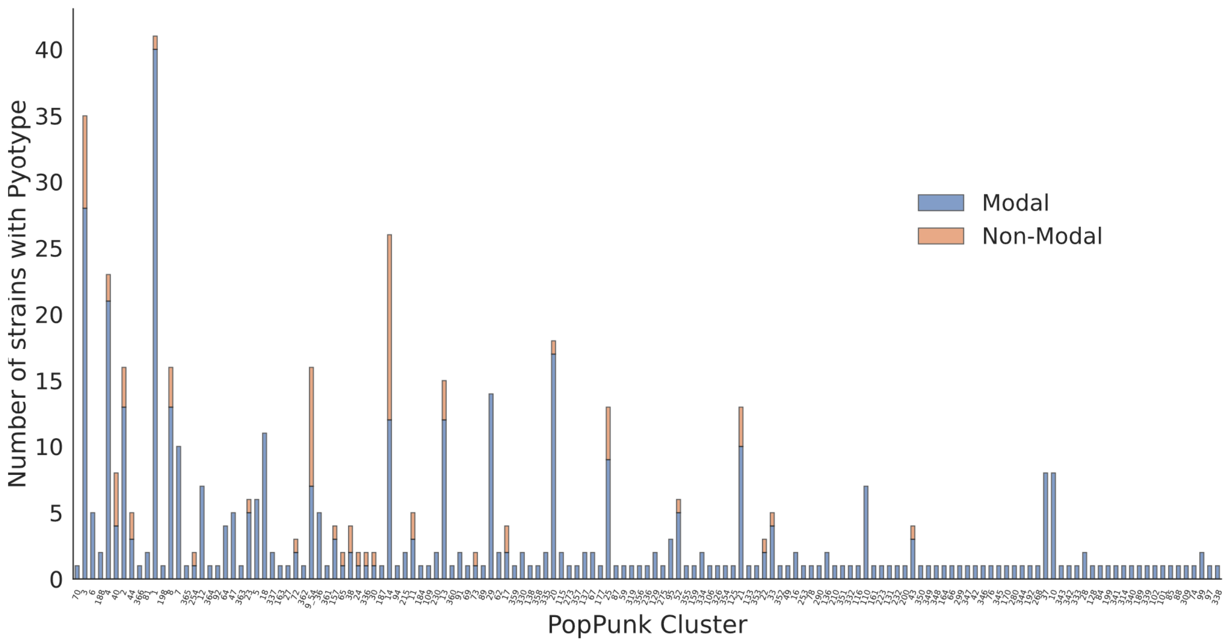

Supplementary Figure 3: Number of strains within each poppunk cluster that contain the modal pyocin type (the most common combination of pyocins and immunity proteins).

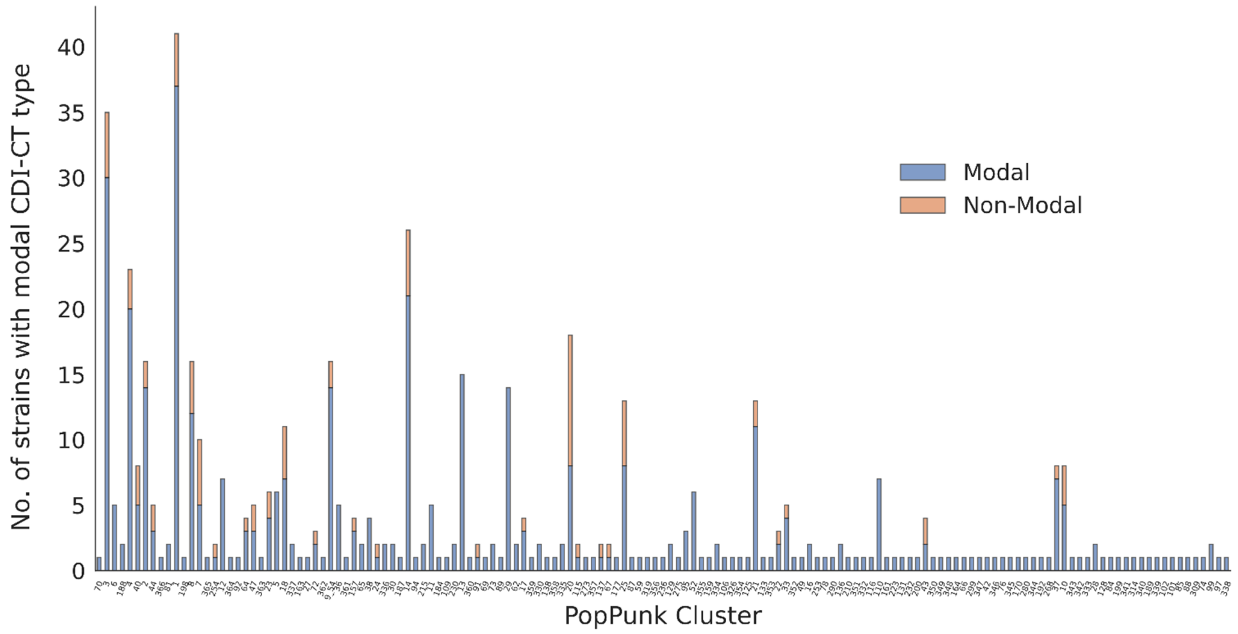

Supplementary Figure 4: Number of strains within each poppunk cluster that contain the modal CDI-CT type (the most common combination of CDI cytotoxic domains).

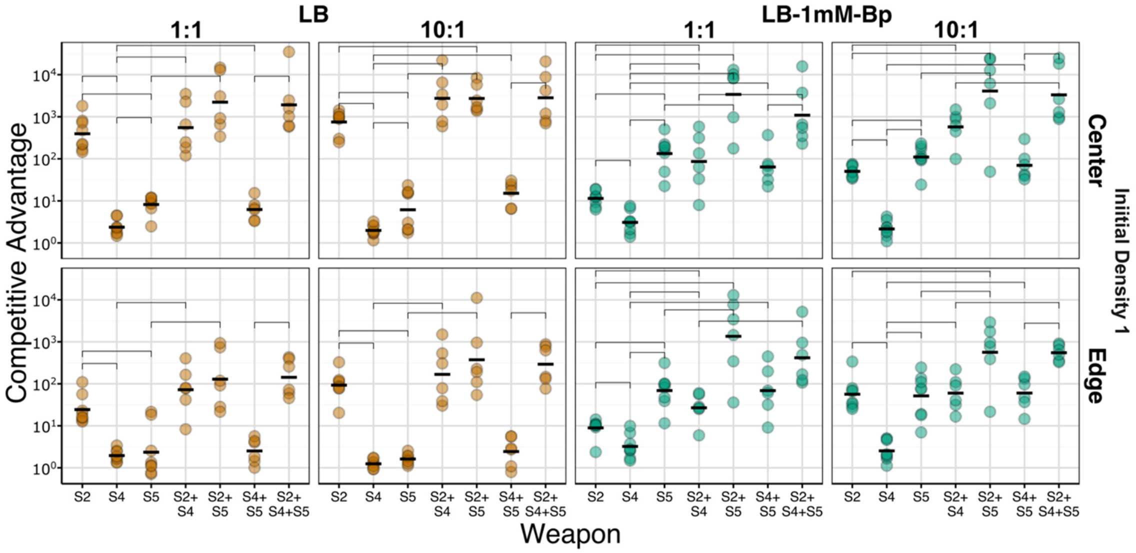

Supplementary Figure 5: Competitive advantage provided by single and combinations of the S pyocins of *P. aeruginosa* PAO1 in colony biofilms. Mutant strains had either single or combinations of deletions of both the toxin and immunity protein of pyocins S2, S4, and/or S5, and were tagged with Tn7-GmR-mScarlet/eYFP. Competitions were carried out on either LB 1.5% agar (orange) or LB 1.5% agar + 1mM bipyridyl (Bp) (blue), inoculated at an initial density of OD600 1.0 and an initial ratio of either 1:1 or 10:1 wild-type to pyocin susceptible. Competitive advantage was determined by spot-plating and counting numbers of CFU. Brackets indicate significant differences ($p < 0.05$) according to BH-corrected pairwise t-tests.

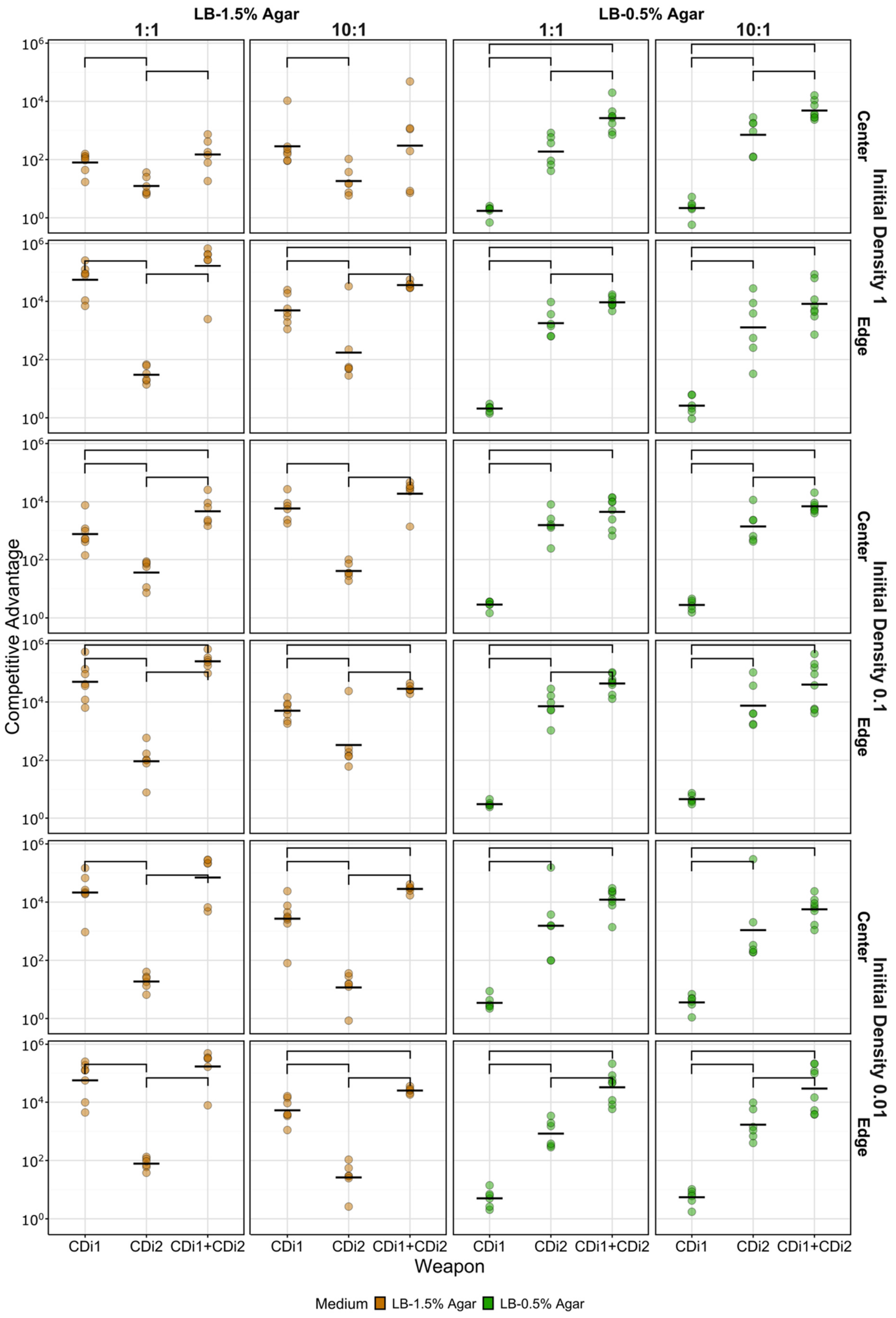

Supplementary Figure 6: Competitive advantage provided by single and combinations of CDI systems of *P. aeruginosa* PAO1 in colony biofilms. Mutant strains had either single or combinations of deletions of both the toxin and immunity protein of CDI1 and/or CDI2, and were tagged with Tn7-GmR-mScarlet/eYFP. Competitions were carried out on either LB 1.5% agar (orange) or LB 0.5% agar (green), inoculated at an initial density of OD600 1.0, 0.1, or 0.01, and an initial ratio of either 1:1 or 10:1 wild-type to CDI susceptible. Competitive advantage was determined by spot-plating and counting numbers of CFU. Brackets indicate significant differences ($p < 0.05$) according to BH-corrected pairwise t-tests.

**Table S1**: Outer membrane pyocin receptors and the seed sequences used to identify them with abricate, Pfam domains used to identify pyocin cytotoxic domains and associated immunity proteins[12], and Pfam domains used to identify CDiI toxins (cdiA1/2) or transporters (cdiB). * indicated a pHMM generated for this study and can be found in Supplementary data.

| **Outer membrane pyocin receptor** | **Accession** | **Pyocin(s) which target receptor** | |
|---|---|---|---|
| FpvAI | EU348646.1 | S2, S4, S7, S11 | |
| FpvAIIa | AY765263.1 | S3, S12 | |
| FpvAIIb | AF540992.1 | S3, S12 | |
| FpvAIII | AY765261.1 | - | |
| FptA | PA4221 | S5 | |
| fiuA | PA0470 | M1 | |
| hxuA | PA1302 | S1, S13, M4 | |
| **Cytotoxic domain** | **Pyocins** | **Cytoxic domain Profile (Pfam)** | **Immunity Domain Profile** |
| HNH-DNase | S1, S2, S8, AP41, S9 | PF12639.2 | PF01320.13 |
| Non-HNH DNase | S3 | *S3DNase | *S3DNase_imm |
| S4-like tRNase | S4 | PF12106.3 | PF11480.3 |
| ColE3-like tRNase | S6, S7 | PF09000.5 | E3-like_imm [37] |
| Pore forming | S5 | PF01024.14 | PF03526.8 |
| Lipid II degradation | M1, M4 | PF14859.1 | - |
| ColD-like tRNase | S11,S12,S13 | PF11429.3 | PF09204.5 |
| **Pfam domain** | **Accession** | **cdiA/B** | |
| Fil_haemagg | PF05594.9 | cdiA | |
| Fil_haemagg_2 | PF13332.1 | cdiA | |
| Haemagg_act | PF05860.8 | cdiA | |
| POTRA_2 | PF08479.6 | cdiB | |
| ShlB | PF03865 | cdiB | |

**Table S2:** Strains of *Pseudomonas aeruginosa* PAO1 and *E. coli* constructed and used in this study.

| **Strain** | **Genotype** | **Purpose** |
|---|---|---|
| WT PAO1 | Wild-type | Attacker in CDI competitions |
| ΔS2 | Deletion of PA0985 (S5) and PA0984 (S5I) | Susceptible in pyocin competitions |
| ΔS4 | Deletion of PA0985 (S5) and PA0984 (S5I) | Susceptible in pyocin competitions |
| ΔS5 | Deletion of PA0985 (S5) and PA0984 (S5I) | Susceptible in pyocin competitions |
| ΔS2 ΔS4 | Deletion of S2/S2I and S4/S4I | Susceptible in pyocin competitions |
| ΔS2 ΔS5 | Deletion of S2/S2I and S5/S5I | Susceptible in pyocin competitions |
| ΔS4 ΔS5 | Deletion of S4/S4I and S5/S5I | Susceptible in pyocin competitions |

| | | |
|---|---|---|
| ΔS2 ΔS4 ΔS5 | Deletion of S2/S2I, S4/S4I, and S5/S5I | Susceptible in pyocin competitions |
| ΔCDI1 | Deletion of CDI1 and CDI1I (PA0040-PA0041) | Susceptible in CDI competitions |
| ΔCDI2 | Deletion of CDI2 and CDI2 I (PA2462-downstream ORF) | Susceptible in CDI competitions |
| ΔCDI1 ΔCDI2 | Deletion of CDI1/1I and CDI2/2I | Susceptible in CDI competitions |
| *E. coli* JKE201 pOPC-244 | Conjugation of pOPC-244 (fpvA::YPet, pilM::mCherry, GmR) | Delivery of plasmid into PAO1 |
| *E. coli* JKE201 pOPC-246 | Conjugation of pOPC-246 (fpvA::YPet, pilM::mCherry, GmR) | Delivery of plasmid into PAO1 |
| PAO1 pOPC-244 | PAO1 wt carrying plasmid | Measurement of receptor expression |
| PAO1 pOPC-246 | PAO1 wt carrying plasmid | Measurement of receptor expression |

**Table S3**: Primers used for construction of deletion mutants.

| Primer | Sequence |
|---|---|
| S2-out-F | ggctttattcgtcgaacaatggc |
| S2-out-R | cttgaccttacgagtcatccgc |
| S2-del-UpF | caagcttctgcaggtcgactctagaggatccgtatatcggccaggacttcaagc |
| S2-del-UpR | ggttcgtaatcattgacagccataga |
| S2-del-DownF | ccacaagggagggaagtgatatg |
| S2-del-DownR | cccgtggaaattaattaaggtaccgaattcgcaacgatgtgttctcgaatcc |
| S2I-del-Down-F | ctttaaggccggttagttggcc |
| S2I-del-Down-R | gcaacgatgtgttctcgaatcc |
| S4-out-F | agtaccgaagggtgcctgag |
| S4-out-R | ccttagattggcgagaagcgc |
| S4-del-UpF | ggcgcactattatttgtcatgaatgg |
| S4-del-UpR | ggcagtgtcttcgaacactcg |
| S4-del-DownF | tgtgtagcggcttgcaatcg |
| S4-del-DownR | cgggcgaatcatctggagaaataag |
| S4I-del-DownF | tgtgtagcggcttgcaatcg |
| S4I-del-DownR | aaagccattcatgacaaataatagtgcgccctgagttggcttgcaatcccttg |
| S5-out-F | tagaggctgtggactacccc |
| S5-out-R | tcggcaaactcagacgttcg |
| S5-del-UpF | ggccttgcttggttttaaatggatattacgttgtcattggacatttagacttctcc |
| S5-del-UpR | cccgtggaaattaattaaggtaccgaattcttcaccatcaagccggcag |
| S5-del-DownF | aagcttctgcaggtcgactctagaggatcctacgtcgatgatctgggccatc |
| S5-del-DownR | aatggagaagtctaaatgtccaatgacaacgtaatatccatttaaaaccaagcaaggcc |
| S5I-del-UpF | aagcttctgcaggtcgactctagaggatccgctttgagcataggaggaagtcc |
| S5I-del-UpR | cattaaaccctacgaggcctttccatacttgtatttaaagctcaattagcaccccg |
| S5I-del-DownF | cggggtgctaattgagctttaaatactattggaaaggcctcgtagggtttaatg |
| S5I-del-DownR | cccgtggaaattaattaaggtaccgaattcgctttgcctccattggtcagtc |
| CDI2-out-F | gaaaataaatccgcccccttttatgc |
| CDI2-out-R | catcgagaccggcaagaacc |
| CDI2-del-UpF | aagcttctgcaggtcgactctagaggatcctgaagcagcgtagactctagcc |
| CDI2-del-UpR | cccgtggaaattaattaaggtaccgaattccagcgcaccaacaactatctcg |
| CDI2-del-DownF | aagcttctgcaggtcgactctagaggatcctagccatggaaagatctgcccc |
| CDI2-del-DownR | ttgaagtcgtagtttttaaaacatacatcg |
| CDI2I-del-DownF | ttgaagtcgtagtttttaaaacatacatcggctagaacgcaacatccgatgtc |
| CDI2I-del-DownR | cccgtggaaattaattaaggtaccgaattccagcgcaccaacaactatctcg |
